# MammalMethylClock R package: software for DNA Methylation-Based Epigenetic Clocks in Mammals

**DOI:** 10.1101/2023.09.06.556506

**Authors:** Joseph A. Zoller, Steve Horvath

## Abstract

Epigenetic clocks are prediction methods that relate cytosine methylation levels at specific genomic sites to chronological age, accounting for nuances such as genome mapping information and possibly life history traits such as age at sexual maturity. Defined as multivariate regression models, these DNA methylation-based biomarkers have gained prominence in species conservation efforts and preclinical studies. Once built, these models predict an individual’s age from their tissue sample’s methylation profile, with great potential for aging research, disease risk prediction, gauging intervention impacts. Through the collaborative efforts of our Mammalian Methylation Consortium, we’ve described epigenetic clocks for numerous mammalian species. Our R package includes the utility for implementing these pre-existing mammalian clocks. Overall, we present software tools and a dedicated R package called *MammalMethylClock*, designed for the construction and assessment of epigenetic clocks.

## INTRODUCTION

Aging is intertwined with many molecular modifications ^1^. Among these, cytosine methylation is particularly notable for its ability to facilitate the creation of pan-tissue aging clocks—age estimators based on multivariate regression models that are applicable to all tissues ^2–5^. The initial human epigenetic clocks were designed leveraging the capabilities of the Illumina Infinium methylation array platform ^2,6,7^. This was followed by the advent of mouse methylation clocks, developed using a different sequencing-based platform ^8–10^. Our Mammalian Methylation Consortium employed the mammalian methylation array which lends itself for measuring methylation levels in upwards of 36,000 CpGs, especially those flanking DNA sequences that are conserved across mammals ^11^. Through this initiative, we successfully profiled an astounding 11,754 samples. These spanned 59 tissue types and were sourced from 185 mammalian species representing 19 taxonomic orders. Remarkably, the age range of these samples extends from prenatal stages right up to a venerable 139 years, as seen in the bowhead whale (Balaena mysticetus) ^12,13^. We published numerous epigenetic clocks tailored to specific species and tissue types ^14–38^. The main objective of this article is to present the R software tools and the accompanying statistical techniques pivotal for the development, assessment, and application of these epigenetic clocks.

In the following, we will describe the steps to use this software package, including how to apply the published non-human and multi-species clocks, and the Universal Pan-Mammalian clocks ^13^, to user generated mammalian methylation data of the same species, and how to build new clocks based on user generated mammalian methylation training data.

## RESULTS

### Overview of the software package

The MammalMethylClock R package serves as a comprehensive suite with various components tailored to the demands of constructing, assessing, and deploying new epigenetic clocks. An overview of the typical analysis steps can be found in **Figure 1**.

**Figure 1.**
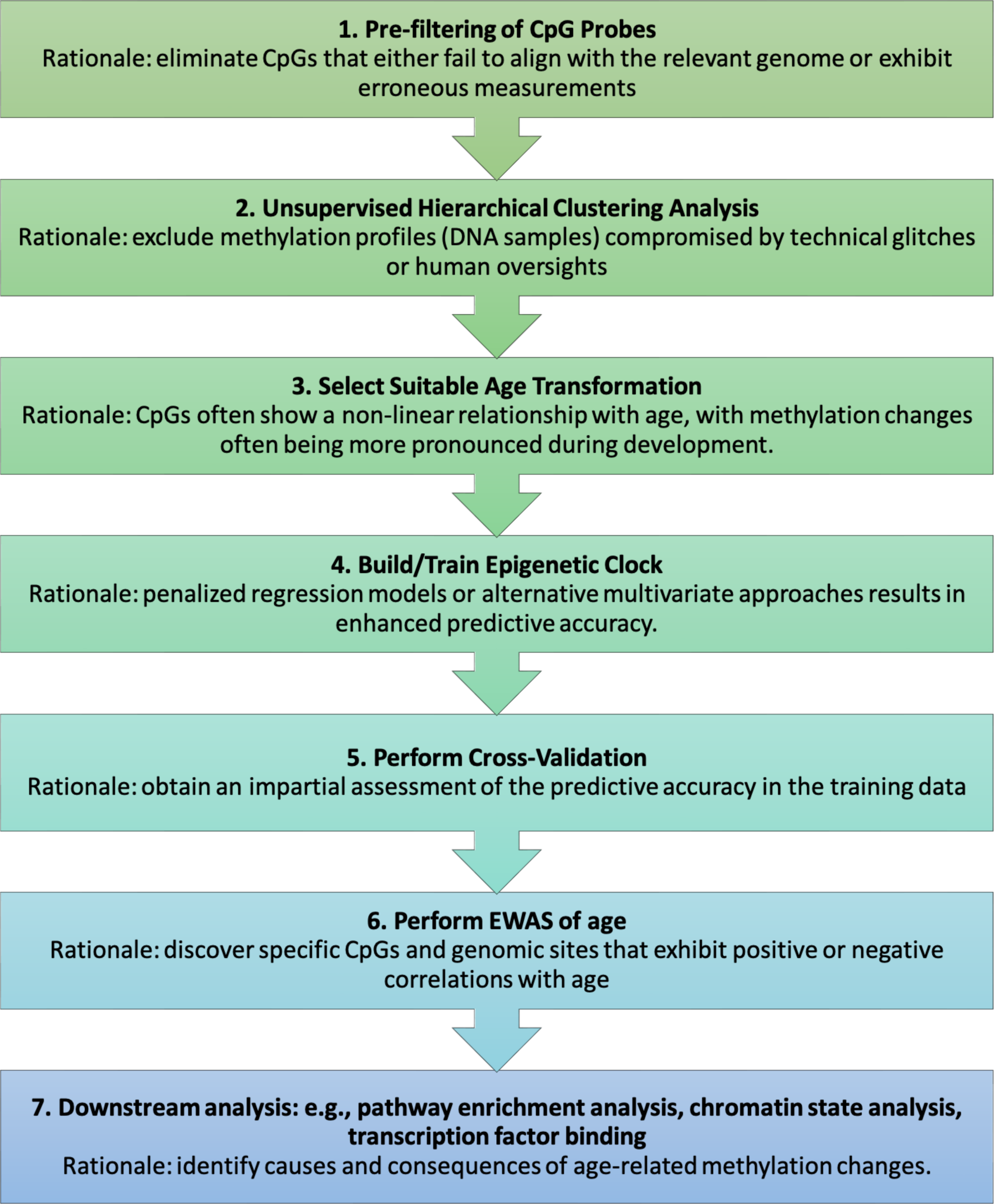
Flowchart for Constructing and Evaluating Epigenetic Clocks. This flowchart outlines the foundational stages of creating and appraising epigenetic clocks, alongside the typical procedures and their underlying logic. For a deeper understanding of the causes and effects of epigenetic clocks, one might employ functional enrichment tools and various other computational resources.

To commence, the **Building Clocks** module empowers users to fashion new epigenetic clocks. This is facilitated using training data supplied by the user, which should encompass precise age values (outcome variable) alongside normalized methylation data (covariates also known as features).

Our package also offers an **Applying Clocks** function, designed to predict the DNA-methylation age for a (test) data set. To achieve this, users need to provide normalized methylation data and their chosen epigenetic clock, which typically comprises a coefficients table. Additionally, if pertinent, an inverse age transformation should also be incorporated.

Recognizing the criticality of rigorous model evaluation, the package integrates two distinct cross-validation schemes: **Leave-One-Out (LOO)** and **Leave-One-Species-Out (LOSO)**. The LOO strategy adopts an approach where each sample undergoes evaluation by being singularly excluded. In contrast, the LOSO method operates on a broader spectrum, whereby every batch of samples stemming from an individual species is set aside once for the assessment.

Intriguingly, for researchers keen on probing the relationship between DNA methylation and age, the **Epigenome-Wide Association Studies (EWAS)** feature has been incorporated. Lastly, the package encompasses the **Meta EWAS** utility. This meta-analysis R functions delves deeper into the relationship between DNA methylation at specific CpGs and specific phenotypes, like age, across different tissues or species. However, it introduces an additional layer of nuance by considering a stratifying variable, most commonly the tissue type or species.

### Epigenetic clocks as penalized regression models

Epigenetic clocks, by design, harness statistical models to correlate DNA methylation levels at specified genomic sites with chronological age. This association is further refined with meticulous incorporation and consideration of biological factors, including array probe mapping to the genome and the intrinsic life history traits of the organism. The resultant model offers the potential to predict an individual’s age through the methylation profile derived from a respective tissue sample. This functionality bears significant implications, enabling deeper insights into the aging processes, more accurate disease risk predictions, and the evaluation of various interventions.

Formally, these epigenetic clocks are conceptualized as regression models. They are typically structured as multivariate linear regression models, wherein the dependent variable is a transformed version of chronological age. The mathematical representation is given as:

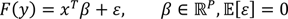

Here, *y* is Age (in years), and *x* ∈ [0,1]^*P*^ signifies a vector of DNA methylation beta values from *P* CpGs. These beta values render a spectrum wherein ‘1’ epitomizes consistent hypermethylation and ‘0’ characterizes consistent hypomethylation. The function *F* serves as the “age transformation” integral to the epigenetic clock, although in numerous applications, it is simply the identity function.

Central to the construction of most epigenetic clocks is a penalized regression model such as elastic net regression or LASSO ^39,40^. As an advanced form of linear regression, elastic net amalgamates the robustness of both LASSO and ridge regularization. This merger empowers the model to select a set of DNA methylation sites (CpGs) from training data sets. These sites, in tandem, are predictive of the outcome variable (e.g., transformed age), and the model estimates their coefficients accordingly. Elastic net regression balances model size with prediction accuracy. It’s adept at fikng a regression model with a limited sample size (e.g., n=50 tissues samples) to high dimensions (37k cytosines) while circumventing overfikng. The regression modeling software employs R functions of the “glmnet” package, serving both the construction and cross-validation analysis of clocks ^41^.

## EWAS

Epigenome-Wide Association Study (EWAS) is used to identify individual CpGs and genomic locations that relate to age. During an EWAS, correlations between every CpG probe and a specified outcome variable (such as age) are ascertained across all (training) data set samples. Pearson correlation test p-values and optionally a local false discovery rate known as q-value can be calculated for each CpG. The EWAS approach allows one to identify CpGs probes with the most significant correlations to the outcome variable. To facilitate this analysis, our software leverages R functions from the “WGCNA” package ^42^ but many other R software tools have been developed for EWAS.

The outcomes of an EWAS act as precursors for further downstream analyses that aim to elucidate the causes or consequences of age-related changes in methylation levels. Examples include the Gene Ontology (GO) enrichment analysis, Genomic Regions Enrichment of Annotations Tool (GREAT) analysis ^43^, and universal chromatin state analysis ^44^. Several special software tools have been developed to identify transcription factors, pathways, chromatin states, and upstream regulators related to age related shifts in DNA methylation. These functionalities are not embedded in our current software package, given that dedicated tools already cater to such downstream analyses.

### Overview of methylation data handling

Our mammalian methylation consortium used the HorvathMammalMethylChip40 Infinium array platform, known as the mammalian methylation array, which profiles oligonucleotide probes that pinpoint sites conserved across a vast majority of mammalian species ^11^. Every published clock integrated into our software package owes its genesis to this mammalian methylation array, a notable example being the Universal Pan-Mammalian clocks ^13^.

Raw array data (idat files resulting from the iScan machine) require normalization to arrive at methylation values (beta values). Our Mammalian Methylation Consortium used the SeSaMe normalization, a tried-and-tested method for all clocks embedded in this software ^45^. The SeSaMe normalization rectifies technical biases. For normalization, we typically use the SeSaMe normalization pipeline but other approaches can be adopted as well ^11^.

Upon normalization, each CpG probe in each sample will manifest a beta value, a fractional representation (ranging between 0 and 1) signifying the methylation extent at that specific CpG site within the given sample.

### CpG probe pre-filtering

For a given array, a histogram of the beta value should typically exhibit a bimodal distribution, with the majority of CpG probes clustering near values of 0 or 1. However, in many species, what we often observe is a trimodal distribution, characterized by an additional intermediate peak around 0.5. This mid-point peak, which is represents a technical artefact, signifies the presence of CpGs on the array that fail to align with the underlying genome. For instance, the mammalian array features CpGs that align exclusively with humans and not with mice.

For enhanced accuracy and efficiency of epigenetic clocks, it’s common to initiate a pre-filtering process for probes. This is crucial as not every probe on an array might exhibit relevance, especially when dealing with non-human mammalian species. The triad of widely accepted pre-filtering methods comprises:

#### Annotation Mappability Filtering

The mammalian array’s CpG probes aren’t universally applicable to every species. While most CpGs are relevant to humans, a several thousand aren’t applicable for mice ^11^. We advise data analysts to exclude CpG probes from their analysis when the associated oligonucleotide sequence isn’t found in the target genome. The vast majority of our epigenetic clocks are based on CpGs that map to the respective genome. Although our R package doesn’t directly contain genome annotation files, CpG genome annotations for many hundreds of species (including non-mammalian species) are available on the Mammalian Methylation Consortium’s GitHub page ^12,13^ (Horvath, S., Lu, A.T., Li, C.Z., Haghani, A., Arneson, A., Ernest, J. & Mammalian, M.C. https://doi.org/10.5281/zenodo.8180547. (2022)) at https://github.com/shorvath/MammalianMethylationConsortium/tree/v2.0.0.

#### Middle Filtering

It can be advisable to exclude CpG probes with an average DNA methylation value approximating 0.5 across multiple samples. This criterion stems from the observation that CpGs failing to align with the core genome often assume values near 0.5. This specific procedure can be accessed within the software package through the “selectProbes.middleFilter” function.

#### Sesame Detection P-value Filtering

Adopting this method means discarding CpG probes with detection p-values exceeding 0.05. These p-values are ascertained during the sesame normalization of the raw methylation data set. Because this filtering method is done using supplemental data generated during the SESAME normalization ^45^.

### Unsupervised hierarchical clustering

To filter out technical outliers and identify inherent batch effects in the data, unsupervised hierarchical clustering is applied to the normalized methylation data. This technique adeptly highlights samples that differ from expected paperns, such as those grouped by species or tissue. Using the inter-array Pearson correlation, we often employ average linkage hierarchical clustering in our studies. Such unsupervised methods help in understanding primary influences on methylation changes, like tissue type. They can efficiently identify samples that stray from expected tissue types, possibly due to human data entry errors or plate map mismatches. Hierarchical clustering further aids in detecting data entry errors, storage or processing-induced batch effects, and individual DNA samples with unsuccessful methylation measurements, often because of insufficient DNA.

Our go-to method for clustering is the standard R function, ‘hclust’ ^46^. Each branch of the hierarchical clustering dendrogram represents a cluster (group of closely correlated DNA samples). These clusters can be identified various branch cukng techniques, with ‘cutreeStatic’ or ‘cutreeDynamic’ from the WGCNA package ^42^. Hierarchical clustering effectively highlights outlier samples based on a large value of the height value (y-axis) of the cluster tree. To evaluate the pairwise resemblance in methylation array readings, we employ the inter-array Pearson correlation coefficient across CpGs. Generally, anticipated pairwise correlations of the same tissue exceed 0.9. When inpukng into ‘hclust’, utilizing ‘1 minus the correlation’ as a dissimilarity measure proves beneficial, as it renders the height parameter in the resultant clustering tree more interpretable. For discerning intergroup disparities, we usually choose “average linkage” but other intergroup dissimilarities can be used as well.

### Transforming age in the clock model

Age transformations play a pivotal role in bolstering the predictive accuracy of epigenetic clocks outside the training data set. These transformations arise from the observed non-linear correlations between DNA methylation and age. By utilizing straighXorward transformations or drawing from species-specific life history traits, age transformations recalibrate the inherent relationship. When leveraging epigenetic clocks, it’s imperative to incorporate the relevant age transformation into the prediction, especially if such a transformation was pivotal in the clock’s original construction. This software suite offers predefined transformation functions (along with their inverse functions) tailored to the specific pre-constructed clock in use.

### Existing clocks included in the package

The MammalMethylClock package is equipped with built-in clocks designed for a diverse range of mammalian species and groups, spanning from rats, primates, and cetaceans to marsupials. In **Figure 2**, we present a sample of these clocks; namely a summary of the accuracy of all primate-based clocks included in this package, based on cross-validation analysis copied from each of the original publications. These integrated clocks enable users to swiftly determine the biological age of their mammalian tissue samples. To ensure accurate results, the samples should be based on normalized DNA methylation data obtained from the Horvath mammalian array, and the “predictAge” function should be employed. It’s imperative to highlight that these clocks are specifically optimized for data derived from the Horvath mammalian array (or data imputed to align with this format) as their training was based on this array.

**Figure 2.**
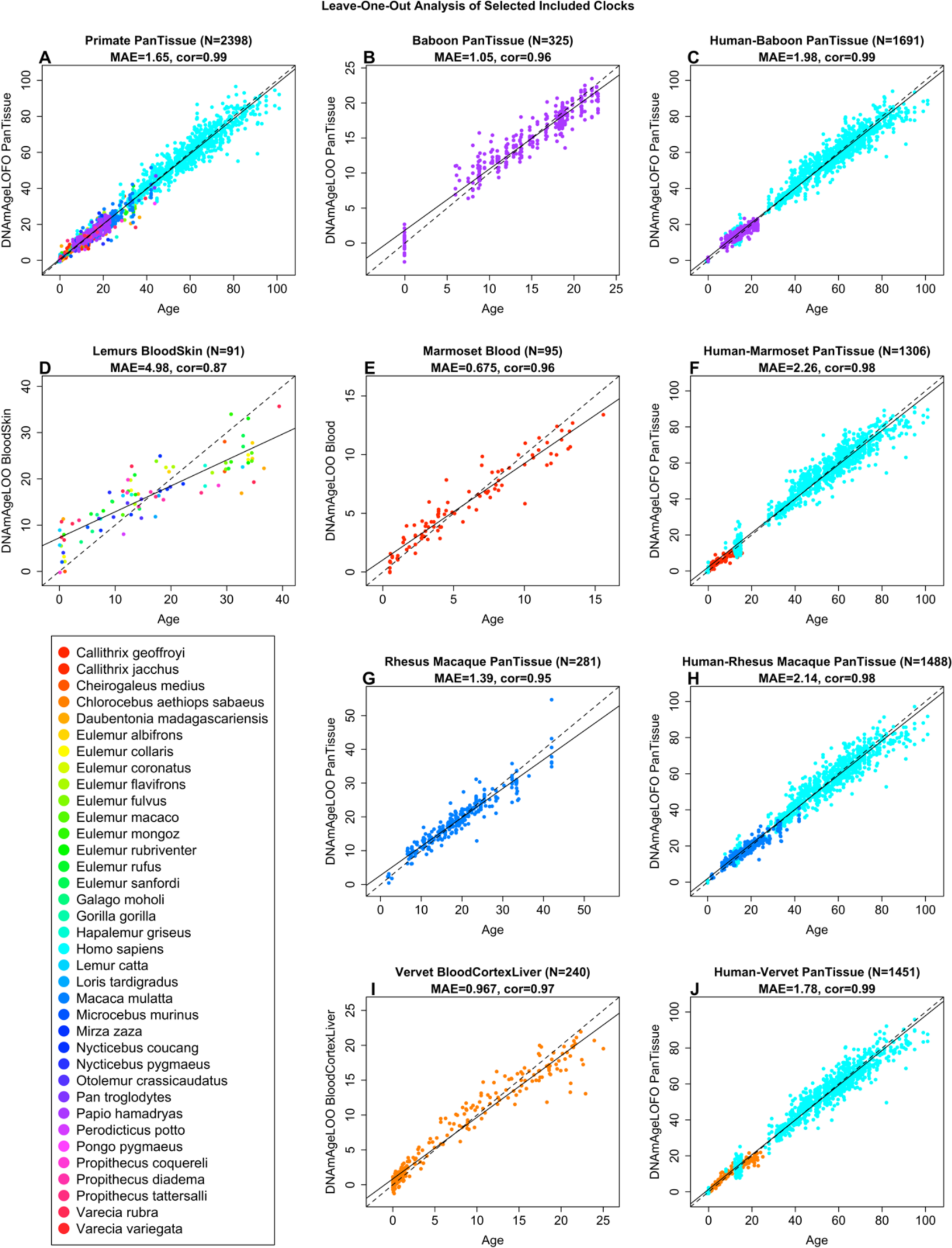
Epigenetic Clocks for Different Primate Species Included in MammalMethylClock. Ten clocks are presented here that are trained on primate species. In order, these clocks are (A) Pan-Primate Pan-Tissue (B) Olive Baboon Pan-Tissue (C) Human-Olive Baboon Pan-Tissue (D) Lemurs Blood-Skin (E) Common Marmoset Blood (F) Human-Common Marmoset Pan-Tissue (G) Rhesus Macaque Pan-Tissue (H) Human-Rhesus Macaque Pan-Tissue (I) Vervet Blood-Cortex-Liver (J) Human-Vervet Pan-Tissue.

To explore the clocks that come pre-installed with this package and obtain crucial details for their appropriate application and citation, utilize the “getClockDatabase()” function.

Additionally, a comprehensive list and other pertinent information can be found on the software’s GitHub repository (https://github.com/jazoller96/mammalian-methyl-clocks/tree/main), and a copy of this list has been directly shared in **Table 1**. For an in-depth guide on leveraging the “predictAge” function, users are directed to the R package documentation accessible within RStudio or the GitHub page. To support ease of access, the coefficient tables for all the included clocks, which are accessible from the individual publications corresponding to each set of clocks, have also been copied in **Supplementary Data 1**.

**Table 1.**
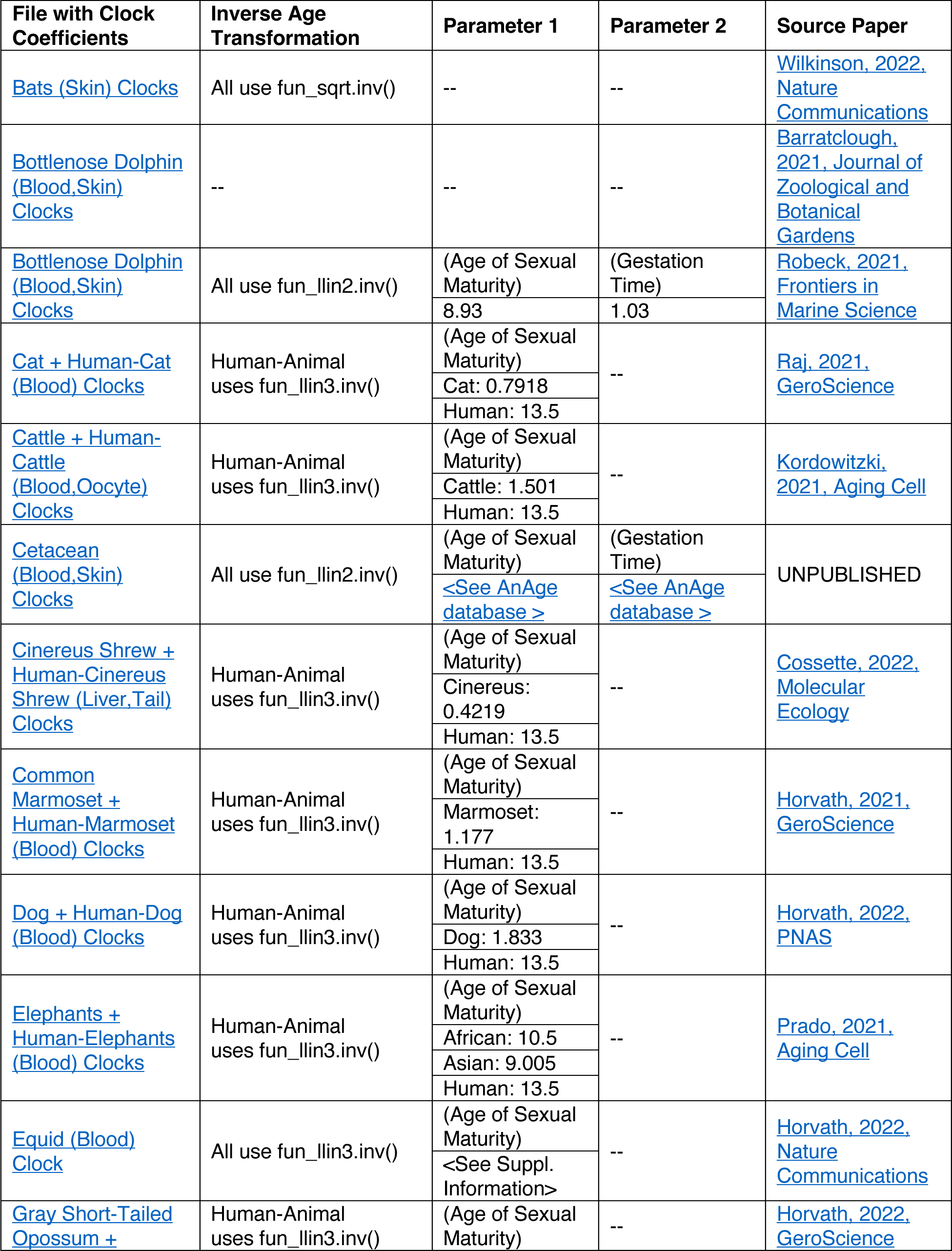
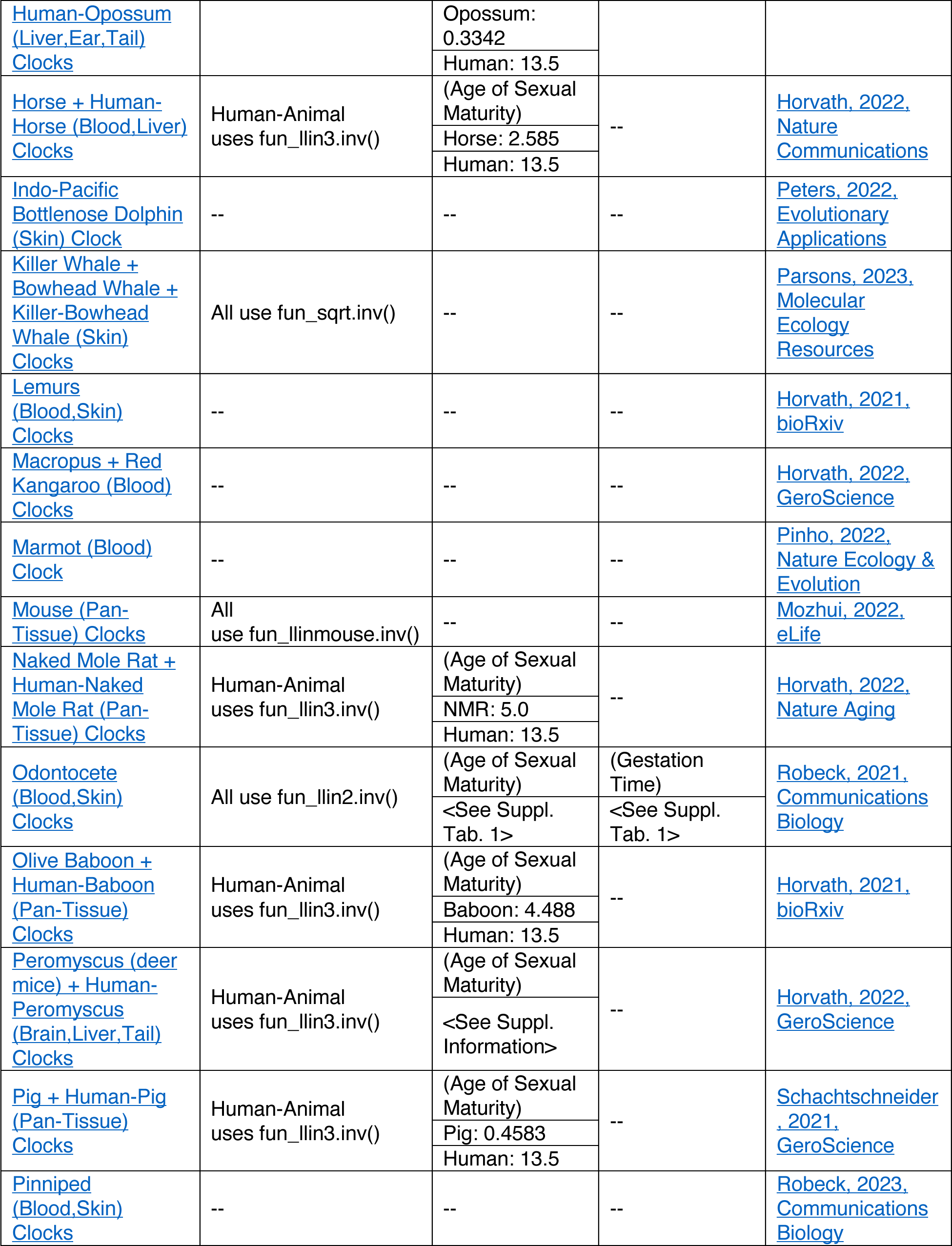
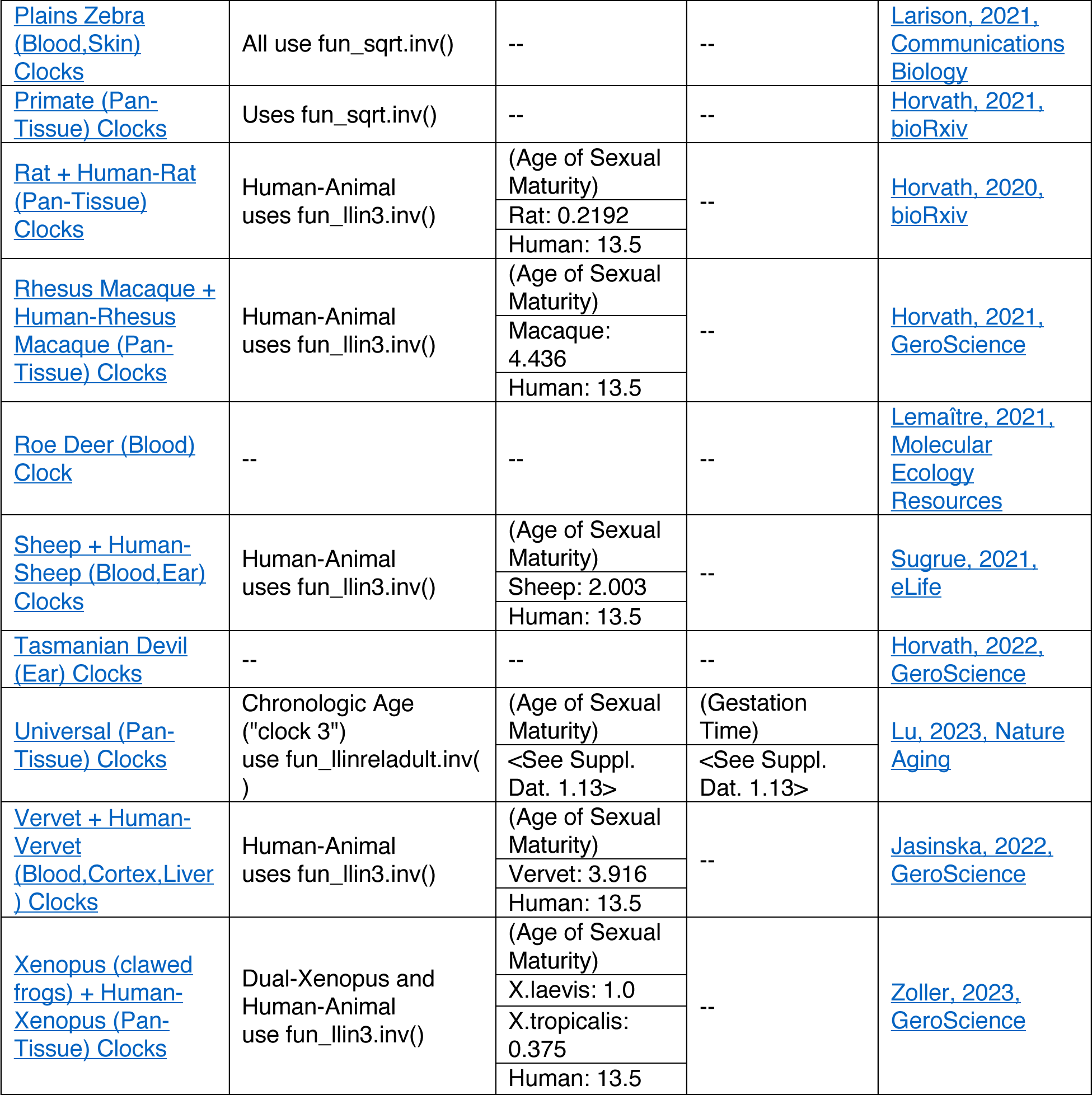
Summary of Existing Clocks Included in MammalMethylClock. This table presents all of the published clocks that are included in the R package, as part of the “predictAge()” function. This table is also provided on the GitHub page for MammalMethylClock, and the online table is updated as new clocks are published and added into the package.

## DISCUSSION

The MammalMethylClock R package offers specialized statistical and bioinformatics tools dedicated to developing epigenetic clocks for single species and multiple species. The R functions can be used to both devise new epigenetic clocks and deploy pre-existing ones. The package includes pre-set clocks developed for many mammalian species analyzed by our Mammalian Methylation Consortium. It caters to a broad research community, accommodating both biologists interested in utilizing existing clocks and computational biologists keen on developing innovative ones.

For constructing precise clocks, it’s essential for users to consider various age transformations, given the non-linear associations between DNA methylation levels and age. Our software offers a suite of age transformations, informed by our previous research, that cater to these complexities. We also suggest investigating the different penalized regression models available in the glmnet R package. Prioritizing pre-filtering methods for CpG probes is crucial. Users can pinpoint high-quality CpG probes chiefly via CpG annotation mappability and detection p-value filtering. This step becomes even more pertinent in studies involving non-human mammals, as some CpG probes may not be detectable. While our primary focus has been on mammalian species, we’ve recently extended the software’s application to amphibians, like *Xenopus* frogs ^47^. While the MammalMethylClock package is optimized for data from the mammalian methylation array, most R functions apply to data from other measurement platforms as well. As the field of epigenetic clocks progresses, the MammalMethylClock R package stands poised to empower researchers in crafting the next generation of epigenetic clocks.

## METHODS

### Training, Validation, and Downstream Analysis Pipelines

The MammalMethylClock package delivers a comprehensive toolkit for researchers venturing into the domain of epigenetic clock studies. The MammalMethylClock package is designed to incorporate a suite of functionalities tailored for the development, assessment, and utilization of epigenetic clocks. Among its main features is the ability to create new epigenetic clocks or other biomarkers using training datasets provided by users. These datasets should include age values combined with normalized methylation data. It’s worth noting that the outcome of this feature might be influenced by inherent randomness. To ensure reproducibility, users are advised to integrate the “set.seed()” command in their R scripts before employing this function. Another primary feature of the MammalMethylClock package is the capacity to predict DNA-methylation age based on specific samples. This feature utilizes normalized methylation data, coupled with any chosen epigenetic clock, which is characterized by a coefficients table and, if deemed necessary, an inverse age transformation.

For model evaluation, the package offers the Leave-One-Out (LOO) cross-validation technique. In this method, each sample is systematically excluded to assess the accuracy and robustness of the model. As with the clock construction, outcomes from this methodology might exhibit some randomness. Thus, the insertion of the “set.seed()” command prior to its activation is recommended. Furthermore, the package includes a variant of this approach, termed Leave-One-Species-Out (LOSO) cross-validation. Here, instead of single samples, all data from a distinct species is left out during the model evaluation phase. Consistency and reproducibility in results can be maintained with the use of the “set.seed()” command.

Beyond these features, the MammalMethylClock package is equipped to conduct Epigenome-Wide Association Studies (EWAS). This allows researchers to probe the relationship between DNA methylation paperns and specific phenotypes, with a predominant emphasis on age. Complementing this, the package also supports Meta EWAS, which refines the evaluation by taking stratifying variables such as tissue type or species into account.

### Implementing Cross-validation Techniques

To ascertain the accuracy of a clock, we advocate that users delve into cross-validation statistics. The software package boasts two distinct methodologies which hinge on the nature of the data at hand: LOO (Leave-One-Out) and LOSO (Leave-One-Species-Out). Each approach partitions the dataset in a unique manner for validation purposes.

In the LOO analysis, employed via the “saveLOOEstimation” function, the process unfolds for every sample within the dataset as follows: Initially, a provisional replica of the datasheet and associated methylation data is constructed, deliberately excluding the chosen sample from both. Once this is achieved, a Clock model is fit utilizing this provisional dataset. The specifications for this are parallel to those presented in the “saveBuildClock” function. Following this, the coefficient table of this LOO Clock is archived to a list of matrices. Concluding the process, this specific LOO Clock is applied to the omiped sample, and the forecasted value is stored in the output.

For computational efficiency, it’s crucial to understand that this function avoids conducting an internal cross-validation at each iteration to pinpoint a potentially varied optimal value for the penalty parameter lambda for every LOO Clock. Instead, recognizing that these optimal lambda values typically converge closely, the function designates a singular optimal lambda value based on the comprehensive dataset. This identical lambda value is then harnessed to configure every LOO Clock.

### Species characteristics from AnAge

Characteristics of species, including maximum lifespan, age of reaching sexual maturity, and gestational duration, which serve as parameters for certain Clocks within this package, were sourced from a revised edition of the Animal Aging and Longevity Database (AnAge, https://genomics.senescence.info/species/index.html) ^48^. We employ the same enhanced version of AnAge that was presented in our “Universal DNA methylation age across mammalian tissues” ^13^. To support reproducibility, we’ve made available a copy of this table in **Supplementary Data 2**.

## URLs

AnAge, https://genomics.senescence.info/species/index.html

## DATA AND CODE AVAILABILITY

The source code and documentation manual for MammalMethylClock, and clock coefficient .csv files that are included within this software package, can be found on GitHub at https://github.com/jazoller96/mammalian-methyl-clocks/tree/main.

