## Supplementary material for "MammalMethylClock R package: software for DNA Methylation-Based Epigenetic Clocks in Mammals": Table 1

| File with Clock Coefficients | Inverse Age Transformation | Parameter 1 | Parameter 2 | Source Paper |
|---|---|---|---|---|
| Bats (Skin) Clocks | All use fun_sqrt.inv() | -- | -- | Wilkinson, 2022, Nature Communications |
| Bottlenose Dolphin (Blood,Skin) Clocks | -- | -- | -- | Barratclough, 2021, Journal of Zoological and Botanical Gardens |
| Bottlenose Dolphin (Blood,Skin) Clocks | All use fun_llin2.inv() | (Age of Sexual Maturity)<br>8.93 | (Gestation Time)<br>1.03 | Robeck, 2021, Frontiers in Marine Science |
| Cat + Human-Cat (Blood) Clocks | Human-Animal uses fun_llin3.inv() | (Age of Sexual Maturity)<br>Cat: 0.7918<br>Human: 13.5 | -- | Raj, 2021, GeroScience |
| Cattle + Human-Cattle (Blood,Oocyte) Clocks | Human-Animal uses fun_llin3.inv() | (Age of Sexual Maturity)<br>Cattle: 1.501<br>Human: 13.5 | -- | Kordowitzki, 2021, Aging Cell |
| Cetacean (Blood,Skin) Clocks | All use fun_llin2.inv() | (Age of Sexual Maturity)<br><See AnAge database > | (Gestation Time)<br><See AnAge database > | UNPUBLISHED |
| Cinereus Shrew + Human-Cinereus Shrew (Liver,Tail) Clocks | Human-Animal uses fun_llin3.inv() | (Age of Sexual Maturity)<br>Cinereus: 0.4219<br>Human: 13.5 | -- | Cossette, 2022, Molecular Ecology |
| Common Marmoset + Human-Marmoset (Blood) Clocks | Human-Animal uses fun_llin3.inv() | (Age of Sexual Maturity)<br>Marmoset: 1.177<br>Human: 13.5 | -- | Horvath, 2021, GeroScience |
| Dog + Human-Dog (Blood) Clocks | Human-Animal uses fun_llin3.inv() | (Age of Sexual Maturity)<br>Dog: 1.833<br>Human: 13.5 | -- | Horvath, 2022, PNAS |
| Elephants + Human-Elephants (Blood) Clocks | Human-Animal uses fun_llin3.inv() | (Age of Sexual Maturity)<br>African: 10.5<br>Asian: 9.005<br>Human: 13.5 | -- | Prado, 2021, Aging Cell |
| Equid (Blood) Clock | All use fun_llin3.inv() | (Age of Sexual Maturity)<br><See Suppl. Information> | -- | Horvath, 2022, Nature Communications |
| Gray Short-Tailed Opossum + Human-Opossum (Liver,Ear,Tail) Clocks | Human-Animal uses fun_llin3.inv() | (Age of Sexual Maturity)<br>Opossum: 0.3342<br>Human: 13.5 | -- | Horvath, 2022, GeroScience |

| | | | | |
|---|---|---|---|---|
| Horse + Human-Horse (Blood,Liver) Clocks | Human-Animal uses fun_llin3.inv() | (Age of Sexual Maturity)<br>Horse: 2.585<br>Human: 13.5 | -- | Horvath, 2022, Nature Communications |
| Indo-Pacific Bottlenose Dolphin (Skin) Clock | -- | -- | -- | Peters, 2022, Evolutionary Applications |
| Killer Whale + Bowhead Whale + Killer-Bowhead Whale (Skin) Clocks | All use fun_sqrt.inv() | -- | -- | Parsons, 2023, Molecular Ecology Resources |
| Lemurs (Blood,Skin) Clocks | -- | -- | -- | Horvath, 2021, bioRxiv |
| Macropus + Red Kangaroo (Blood) Clocks | -- | -- | -- | Horvath, 2022, GeroScience |
| Marmot (Blood) Clock | -- | -- | -- | Pinho, 2022, Nature Ecology & Evolution |
| Mouse (Pan-Tissue) Clocks | All use fun_llinmouse.inv() | -- | -- | Mozhui, 2022, eLife |
| Naked Mole Rat + Human-Naked Mole Rat (Pan-Tissue) Clocks | Human-Animal uses fun_llin3.inv() | (Age of Sexual Maturity)<br>NMR: 5.0<br>Human: 13.5 | -- | Horvath, 2022, Nature Aging |
| Odontocete (Blood,Skin) Clocks | All use fun_llin2.inv() | (Age of Sexual Maturity)<br><See Suppl. Tab. 1> | (Gestation Time)<br><See Suppl. Tab. 1> | Robeck, 2021, Communications Biology |
| Olive Baboon + Human-Baboon (Pan-Tissue) Clocks | Human-Animal uses fun_llin3.inv() | (Age of Sexual Maturity)<br>Baboon: 4.488<br>Human: 13.5 | -- | Horvath, 2021, bioRxiv |
| Peromyscus (deer mice) + Human-Peromyscus (Brain,Liver,Tail) Clocks | Human-Animal uses fun_llin3.inv() | (Age of Sexual Maturity)<br><See Suppl. Information> | -- | Horvath, 2022, GeroScience |
| Pig + Human-Pig (Pan-Tissue) Clocks | Human-Animal uses fun_llin3.inv() | (Age of Sexual Maturity)<br>Pig: 0.4583<br>Human: 13.5 | -- | Schachtschneider, 2021, GeroScience |
| Pinniped (Blood,Skin) Clocks | -- | -- | -- | Robeck, 2023, Communications Biology |
| Plains Zebra (Blood,Skin) Clocks | All use fun_sqrt.inv() | -- | -- | Larison, 2021, Communications Biology |
| Primate (Pan-Tissue) Clocks | Uses fun_sqrt.inv() | -- | -- | Horvath, 2021, bioRxiv |
| Rat + Human-Rat (Pan-Tissue) Clocks | Human-Animal uses fun_llin3.inv() | (Age of Sexual Maturity)<br>Rat: 0.2192 | -- | Horvath, 2020, bioRxiv |

| | | | | |
|---|---|---|---|---|
| | | Human: 13.5 | | |
| Rhesus Macaque + Human-Rhesus Macaque (Pan-Tissue) Clocks | Human-Animal uses fun_llin3.inv() | (Age of Sexual Maturity)<br>Macaque: 4.436<br>Human: 13.5 | -- | Horvath, 2021, GeroScience |
| Roe Deer (Blood) Clock | -- | -- | -- | Lemaître, 2021, Molecular Ecology Resources |
| Sheep + Human-Sheep (Blood,Ear) Clocks | Human-Animal uses fun_llin3.inv() | (Age of Sexual Maturity)<br>Sheep: 2.003<br>Human: 13.5 | -- | Sugrue, 2021, eLife |
| Tasmanian Devil (Ear) Clocks | -- | -- | -- | Horvath, 2022, GeroScience |
| Universal (Pan-Tissue) Clocks | Chronologic Age ("clock 3")<br>use fun_llinreladult.inv() | (Age of Sexual Maturity)<br><See Suppl. Dat. 1.13> | (Gestation Time)<br><See Suppl. Dat. 1.13> | Lu, 2023, Nature Aging |
| Vervet + Human-Vervet (Blood,Cortex,Liver) Clocks | Human-Animal uses fun_llin3.inv() | (Age of Sexual Maturity)<br>Vervet: 3.916<br>Human: 13.5 | -- | Jasinska, 2022, GeroScience |
| Xenopus (clawed frogs) + Human-Xenopus (Pan-Tissue) Clocks | Dual-Xenopus and Human-Animal use fun_llin3.inv() | (Age of Sexual Maturity)<br>X.laevis: 1.0<br>X.tropicalis: 0.375<br>Human: 13.5 | -- | Zoller, 2023, GeroScience |
